# Reference-Free Germline Immunoglobulin Allele Discovery from B Cell Receptor Sequencing Data

**DOI:** 10.1101/2023.11.25.568681

**Authors:** Ivana Cvijović, Elizabeth R. Jerison, Stephen R. Quake

## Abstract

Antibodies, or immunoglobulins, are a diverse set of molecules that play a critical role in adaptive immunity. They are generated in a process which begins with the recombination of germline V, D, and J gene segment alleles, and refined by hypermutation of these germline sequences upon antigen exposure. Antibody repertoire analysis often requires the knowledge of the germline V, D, and J alleles to detect hypermutations and understand the phylogenetic relationships of related B cells. However, germline immunoglobulin alleles are remarkably diverse and incompletely annotated, making it necessary to construct personalized databases for every individual. Though several approaches for the detection of germline immunoglobulin variants exist, they often rely on refining existing databases using simplifying assumptions about the relationships of germline alleles in a given organism, or about the form of evolutionary process that shapes antibody repertoires. Here, we present `grmlin`, an alternative computational approach to detecting germline alleles. Our approach exploits two empirical properties of B cell repertoires: the abundance of germline sequences in antibody repertoires and the enormous diversity of antibody sequence space, to detect germline alleles from B cell receptor sequencing data without reliance on a reference database. As such, it is in principle applicable to non-model organisms. We validate this approach by detecting the germline alleles of 11 pairs of twins and show that it achieves equivalent sensitivity and better specificity than previous methods.

## Introduction

Antibodies or immunoglobulins are key components of adaptive immune response. In B cell precursors, complete immunoglobulin (Ig) sequences that encode the variable region of the B cell receptor (BCR) are generated by somatic recombination of germline V, D, and J immunoglobulin segments. In this process, each B cell precursor selects one of the many contiguous V, D, and J genes present at the Ig locus, and recombines them in an error-prone way to create a region of enormous diversity at the junction of the recombined segments. Cells that acquire a functional receptor in this process may also undergo further diversification upon antigen exposure in a process known as affinity maturation, during which B cells undergo rounds of somatic hypermutation and selection.

In recent years, it has been possible to survey the diversity of antibody repertoires through targeted sequencing of BCRs. The interpretation of these data often depends on associating each observed BCR sequence with its germline V, D, and J segments. Knowledge of germline sequences of particularly desirable antibodies (e.g. classes of broadly neutralizing antibodies) is often necessary in rational vaccine design [15]. Moreover, the ability to associate BCRs with their ancestral sequences is often central to studying B cell evolution, as it often facilitates the identification of B cells that are clonally related and makes it possible to calculate the absolute number of mutations that a particular B cell has accumulated.

In recognition of these goals, there now exist databases of known Ig germline gene variants in a number of species. These databases reveal several orders of magnitude of variation in divergence between different Ig germline variants in some species, with many genes having a large number of closely related alleles. However, even in the most extensively studied species, these databases are incomplete, with new germline variants continuously identified as more individuals are sequenced. An additional challenge arises due to the fact that individuals only carry a fraction of all of the genes present in these databases (in humans, the heavy chain locus carries roughly 50 out of the over 250 possible V Genes, 27 of the 35 D, and 6 of the 12 known J genes). Because typical BCR divergences from their germline variants is often substantially larger than the divergence between the different allelic variants present in the database (5-10%, or larger, compared to several nucleotides between alleles), simply associating each allele with the closest variant present in the database often leads to the detection of a large number of alleles that are not present in the true germline [12]. Even in cases in which a small amount of uncertainty in the identity of the germline sequence is permissible, this can lead to the incorrect splitting and merging of clonal antibody lineages, and thus to potential biases in the inferred B cell phylogenies and erroneous conclusions about their evolutionary process.

Motivated by these challenges, a number of methods to infer personalized germline Ig databases from BCR deep sequencing data have been developed [2, 4, 5, 12, 18]. These broadly fall into two categories. The first infers germline sets using a hierarchical clustering of observed BCR sequences, which in principle does not rely on an existing database, as implemented in the `IgDiscover` package [2]. The second category of methods, including those implemented in the widely-used `TIgGER` and `partis` packages, infers a personalized gene database by examining mutational patterns at each site in an alignment of sequences that are attributed to a known gene in the IMGT database, [4, 5, 12, 18]. These approaches rely on the existence of a well-annotated database of Ig germline variants, which are refined by the methods by calling mutations at a small number of sites. Though the different methods vary both in the method of attributing sequences to different genes, as well as in the exact procedure used to call germline mutations at each position, they are based on a underlying model of neutral mutation accumulation in BCR sequence evolution.

The complex patterns of divergence between different Ig germline variants, as well as the unique ways in which B cell receptors evolve during affinity maturation pose significant challenges for both clustering and position-based approaches. In the former case, examining Levenshtein distances between known Ig variants shows that neither the sequences of Ig genes present in the IMGT database, nor those of personalized Ig germline databases form particularly tight clusters [2]. Thus, the task of inferring a complete germline gene set based on hierarchical clustering of observed sequences is in principle difficult. In the latter case, the strong selection experienced by B cells during affinity maturation can lead to large and rapid expansions particular BCRs, accompanied by large numbers of closely related hypermutants, which causes significant departures from a neutral mutation-accumulation model [8]. This limits the statistical power of position-based methods, particularly in cases of genes that significantly differ from known Ig variants [12].

Here we present an alternative approach to inferring personalized germline Ig V and J gene databases from deep BCR sequencing data that does not rely on clustering or on prior knowledge of Ig germline genes. Our approach is inspired by existing methods for denoising amplicon sequencing data in microbial community ecology [13, 17] and builds on the intuition that repeated observation of an identical Ig sequence in B cells descended from different rearrangement events (i.e. belonging to different “lineages”) correspond to either germline sequences, or convergent hypermutation events. We show that a simple model of this process can be used to distinguish sequences arising from convergent hypermutation from true germline sequences, by exploiting the fact that unhypermutated, naive B cell sequences represent substantial fractions of unsorted populations of B cells from multiple tissues [10].

We show that while our approach does not rely on or make use of a database of sequences, it primarily identifies known germline variants from human deep BCR sequencing data, and also novel alleles separated by a small number of base pairs. We validate our approach by comparing the inferred Ig germline databases for 11 pairs of twins [7], and show that it returns consistent calls in identical twins, but not necessarily in fraternal pairs. We compare our inferred databases to those inferred by `IgDiscover`, `partis`, and `TIgGER`, and show that our approach has higher sensitivity than `IgDiscover` and equal or better consistency between identical twins than `TIgGER` and `partis`, while not relying on an existing database.

## Results

### Approach

Our approach is motivated by the intuition that identical templated Ig V gene sequences that are seen in B cells descended from different rearrangement events represent either true germline sequences or the results of convergent hypermutation. To identify such sequences and use them to construct a set of true germline species, we must be in a position to: 1) identify the descendants of different recombination events without reference to the templated V and J regions, 2) distinguish between the convergent hypermutants and true germline alleles among these sequences. In this work, we propose a simple approach to both tasks informed by known statistical features of B cell populations that we demonstrate is self-consistent in BCR repertoires. The BCR repertoires that we observe via deep sequencing are generated through a complex biological process that begins with VDJ recombination. For a particular focal germline V segment, the VDJ recombination process founds a number of independent lineages with different J genes and untemplated regions. Some of these lineages later expand and mutate, creating groups of phylogenetically related sequences (**Fig. 1a**). During repertoire sequencing, a subset of extant VDJ sequences is sampled, amplified and sequenced. Crucially, though sequences belonging to the same lineage are generated via a evolutionary process that explicitly violates independence, sequences belonging to different lineages arise independently in the repertoire. Thus, by counting the number of times the same V segment sequence is seen in different lineages, we can estimate the rate at which it arises independently in a repertoire. Here we show that by ranking V segment sequences by the rate at which they are independently created in a repertoire, we can distinguish germline V gene segments from their hypermutated counterparts and thus identify a personalized set of germline genes.

**Figure 1.**
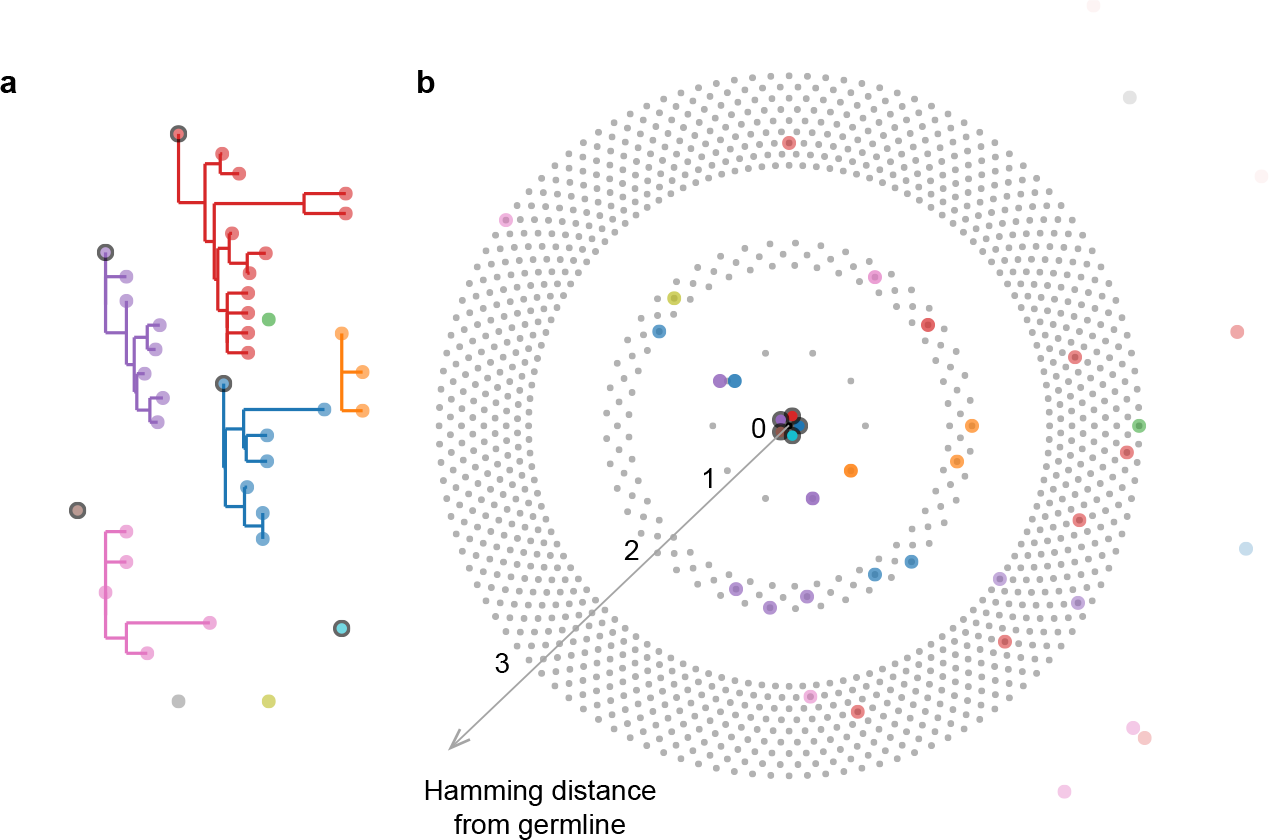
Motivation for approach. (**a**) Schematic depicting the phylogenetic relationships of the antibody transcripts of a sample of B cells that all use the same germline Ig V gene. Points with a black outline represent germline (unhypermutated) sequences. Each color refers to a different lineage, founded by a separate VDJ recombination event. (**b**) Schematic showing occupancy of the possible sequence space by the same sequences, organized by Hamming distance from the germline genes. The first circle represents sequences a single nucleotide apart from the germline sequence, the second sequences two nucleotides from germline, the third sequences three nucleotides apart, and each point represents a possible sequence at that Hamming distance. For visual clarity, this representation refers to hypothetical gene that is 10 nucleotides long, and only has two possible bases at each position, and sequences that are more than three nucleotides apart are not explicitly enumerated by points (in contrast, actual Ig V genes are about 300 nucleotides long, are encoded by four bases, and thus have vastly greater combinatorial complexity). The templated portions of all germline sequences are identical by assumption, independent of the lineage they belong to. As distance from the germline grows, the combinatorial complexity of the space vastly increases. Though sequences that are a small number of nucleotides from the germline may be repeatedly generated in different lineages (see the blue and purple point in the first circle), this becomes increasingly unlikely at greater distances.

Note that, in principle, the ability to rank V segment sequences by the rate at which they are independently created is not alone sufficient for the identification of germline variants: there must also be a separation of scales between the rates of germline sequences (high) and their hypermutated variants variants (low) (**Fig. 1b**). Thus, we need two additional properties in B cell populations. First, unmutated sequences must exist in relatively large numbers. Second, the relative rate of generating any one hypermutated variant must be small compared to the frequency of the unmutated sequence. Neither of these properties need a priori be present. For instance, we would not expect a large proportion of unmutated sequences in sorted memory cell populations. However, naive unhypermutated B cells represent a substantial proportion of all B cells in a range of lymphatic tissues, including the peripheral blood [6]. This means that germline templated sequences are likely to be represented in substantial fractions of BCRs obtained through deep sequencing of unsorted samples of B cells. Second, the templated portions of the variable regions of BCRs are known to harbor a wide diversity of mutations at hundreds of sites (**Fig. 1b**). We demonstrate that though convergently hypermutated sequences do arise routinely within typical B cell repertoires, biases to even the most commonly generated hypermutated sequences are sufficiently weak that they do not outnumber the number of independent lineages carrying germline variants. The combination of these two properties of B cell populations makes it possible to distinguish between germline sequences and convergently hypermutated species derived from these genes simply on the basis of the number of independent occurrences.

### The distribution of mutabilities and the probability of of convergent hypermutation

The probability with which the same hypermutated sequence is created in a lineage will depend on a number of sequence-specific factors, including the fractional mutation rate towards that sequence and the strength of selection for B cells carrying those variants. However, as long as that probability is lower than the fractional abundance of naive cells in the sample, we expect these sequences to represent minority species relative to the germline variant of the same sequence.

The fraction of lineages that contain an unmutated germline variant will depend on a variety of experimental factors, including the age and health status of the subject from which the cells were sampled, tissue of origin and enrichment protocol (e.g. for specific markers) that was applied prior to sequencing. However, in samples consisting of unsorted B cells, these numbers are large: naive B cells represent 50-90% of all B cells in the peripheral blood, lymph nodes, tonsils, and the bone marrow [6]. Moreover, because lineages that contain sequences with the germline variant of a gene tend to be smaller than lineages that do not contain a germline variant, the fraction of sampled lineages that support a germline variant is typically even larger than the fraction of germline cells in the sample (**Fig. S2**).

The probability of convergent hypermutation is more difficult to quantify and depends on a range of processes that occur during B cell development and affinity maturation. These include known known biases in the mutational process that occurs affinity maturation [3, 11, 14, 16, 19], the marginal impact of selection on the templated region during affinity maturation, the dynamics of the affinity maturation process (which may, for instance, deplete weakly hypermutated sequences or make it extremely unlikely to generate variants that are a very large number of mutants away from the mutated sequence), and, lastly, any technical variation in the probability of amplifying and sequencing particular sequences.

However we can make progress by evaluating the strength of these biases in a model-based way for aspects of the process that are better understood, and evaluating the empirical statistics of sequenced BCR repertoires to verify that these are consistent with the assumptions of our overall approach. We begin by examining the biases inherent to the mutational processes that occur during affinity maturation, which have been extensively documented [3, 11, 14, 16, 19]. By examining the inferred distribution of relative mutation rates to different SNVs along the IGHV genes in the IMGT database, we find that though there is substantial variation between the rates at which different SNVs arise, this variation is still moderate enough that the most mutable sites have fractional mutabilities that are uniformly lower than 10% (**Fig. S3**). This implies that convergently hypermutated sequences are expected to be strongly concentrated around germline variants and represented in minority fractions in unsorted samples (**Fig. 1b, Appendix S1**), unless these entropic effects are overcome by selection strongly favoring specific sequences. However, we can show based on crude upper bounds on the effect of selective forces, that these biases do not substantially modify the rate distribution compared to this baseline (**Fig. S4**). We emphasize however that this does not apply to untemplated regions of V and J genes, where nontemplated deletions and insertions during VDJ recombination complicate the process, and certain modifications ultimately end up far likelier than others [3]. Thus, our method cannot reliably infer the presence of germline mutations in the short portions of V genes that may lie within the CDR3.

These observations suggest a simple approach for identifying germline sequences from deep BCR sequencing data, which is to associate the templated portion of each unique Ig V sequence with the number of independent recombination events in which it is observed. Note that this does not involve any clustering of the V sequences, but instead only groups CDR3 segments with similar sequences into the same tentative “lineages” (see **Fig. 2e** and the next section for details). By examining the Levenshtein distances between the sequences associated with multiple recombination events, we can see that they typically form tight, star-shaped clouds consisting of central sequences supported by a very large number of recombination events and surrounded by large numbers of sequences supported by substantially fewer recombination events that are only a few nucleotides apart (see **Fig. 2a**). This picture is consistent with the intuition about the convergent hypermutation process outlined above, which predicts that sequences represented in multiple lineages should form tight clouds of minority species centered around the germline variant.

**Figure 2.**
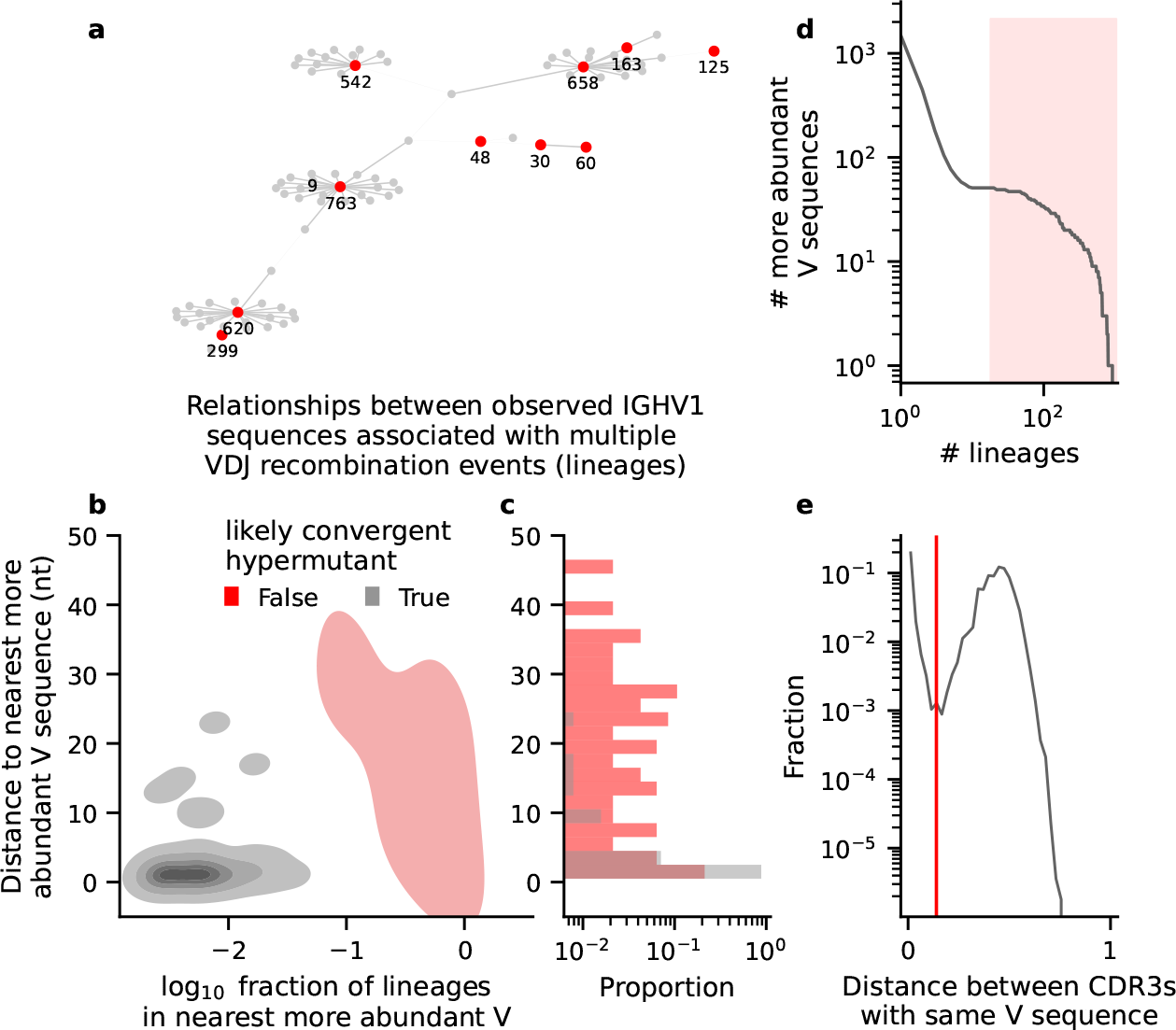
Properties and relationships of recurrently generated IGHV sequences in the human blood. (a)Graph showing the relationships of the observed templated nucleotide V sequences associated with multiple recombination events or lineages in Twin 1A from Ref [7]. The graph was derived from the minimum spanning tree of all V sequences associated with at least 3 lineages and belonging to the IGHV1 family. Edges longer than 3 nucleotides have been eliminated. Red vertices show likely germline V sequences, and grey vertices indicate likely recurrent hypermutants. For vertices supported by more than 5 lineages, the number of lineages supporting them is indicated on the graph. (**b**) For all unique V sequences in the same individual with at least 4 lineages: the distance to nearest more commonly generated nucleotide V sequence in the same IGHV family, and fraction of lineages supporting a V sequence relative to the nearest more commonly hypermutated gene. (**c**) Distribution of distances to the more commonly generated nucleotide V sequence for the same data shown in panel (b). (**d**) Distribution of the number of lineages supporting a unique V sequence across all V families. Red rectangle denotes sequences that account for at least 0.1% of all lineages, which pass the first filter used by `grmlin`. Note the broad plateau in the cumulative density function, which makes the approach relatively insensitive to the exact cutoff implemented. (**e**) Pairwise distances between CDR3s of the same length and associated with the same exact nucleotide V sequence in the same individual. Red line denotes lineage clustering cutoff: CDR3s with smaller distances are considered part of the same lineage and CDR3s at greater distances are considered members of different lineages

Thus we can identify true germline sequences by retaining sequences observed in a sufficiently large fraction of independent recombination events in the dataset (see **Fig. 2d**), and pruning this set of initial candidates by discarding sequences that represent minority species in the vicinity of better-supported alleles (see **Fig. 2a**). By examining the distribution of fractions that these minority sequences represent in BCR repertoires derived from PBMCs drawn from healthy individuals prior to vaccination (previously published in Ref. [7]), we can see that they separate into two prominent groups (**Fig. 2b,c**). The majority of sequences are encountered in fractions smaller than about 5% of the central sequence, consistent with the magnitude of the biases described above. The remainder likely represent other germline alleles, some of which might be very closely related (**Fig. 2b,c**); we retain them by discarding only minority alleles that are represented in fractions consistent with convergent hypermutation. Thus our method in principle cannot resolve very similar germline alleles with differences in usage rates exceeding the inverse of this threshold (in practice, differences of than 10-fold or larger), and similarly to other approaches, it has limited ability to resolve germline variants that are used in fractions larger than rates of convergent hypermutation to distant sequences.

### Implementation

We implemented our approach as a standalone and freely available Python package, which we call `grmlin` (https://github.com/icvijovic/grmlin). The input to `grmlin` is an AIRR-formatted file of nucleotide VDJ sequences sampled from the repertoire, in which the positions of the V gene, the CDR3, and the J gene have been annotated. In cases in which the repertoire belongs to an organism from a species with a known database of possible Ig germline genes, the annotation of the sequence elements can be accomplished with a tool like `IgBLAST` [20]. In non-model organisms, the V gene, J gene, and CDR3 can be parsed by using the known conserved features of immunoglobulin framework (FR) and complementarity determining regions (CDR) [9].

Since each of the Ig V sequences in the repertoire initially represents a candidate germline sequence by assumption, it is crucial that technical artifacts are first removed from the candidate germline repertoire. Specifically, we retain only sequences that contain a CDR3, are in frame, productive, and contain V and J gene fragments of adequate length. In the case of organisms with well-annotated Ig allele databases, we further require that the V and J gene fragments each have at least one high-quality alignment to the database of known alleles. Next, we trim any primer sequences that may be incident within the templated V genes, if relevant to the library design. Finally, we noticed that artifacts associated with base trimming, deletion, or insertion at the very ends of the transcript are common technical artifacts. To address this, we collapse all unique Ig V sequences that are truncations of one another into a single set, and map all the minority members of the set to the most common sequence. Thus, we assume that these minority members likely represent artifacts of the most common sequence; we remove the minority members from further consideration as candidate germline genes, while retaining information about the different lineages they are associated with.

Next, we compile a list of all unique nucleotide V gene sequences, i.e. the 5’ portion of the nucleotide VDJ sequence up to the start of the CDR3. We then annotate each sequence with an approximate number of independent VDJ recombination events, which may each be associated with many closely related CDR3s of the same length. To achieve this, we first compile a list of all unique nucleotide CDR3 sequences associated with that V gene sequence, and then cluster the unique CDR3s of the same length via average linkage clustering. As described above, the clustering condition is defined by the minimum in the nearest-neighbor fractional Hamming distance distribution (see **Fig. 2e**). The total number of CDR3 clusters (of any length) is then used as a proxy for the number of independent VDJ recombination events associated with the unique Ig V nucleotide sequence. As is conventional, we set the default fractional Hamming distance clustering cutoff to 0.15, editable by the user through a command-line parameter.

To be further considered as a germline gene candidate, we require that a V sequence is supported by at least 10 VDJ recombination events, or at least 0.1% of all recombination events observed in the repertoire, whichever is greater (see **Fig. 2d**). This means that our approach inevitably discards very rarely used V genes. Finally, we rank sequences by the number of recombination events supporting them and designate as germline V genes unless they are within a very short Levenshtein distance, *d*_*c*_, from a sequence supported by more recombination events. In the latter case of a sequence in the vicinity of a designated germline V gene, we also designate it as germline if it is supported by a comparable faction of recombination events, *f*_*c*_. We determined *d*_*c*_ and *f*_*c*_ self-consistently by observing patterns of convergent mutation in human data. Specifically, we rarely encounter repeated hypermutation events at distances greater than *d*_*c*_ = 3 edits from the focal germline sequence (**Fig. 2b,c**), and we set *f*_*c*_ to 0.05, as this threshold is higher than the observed hypermutation biases in the repertoire (see **Fig. S4**) for all measured positions at which we have the sufficient power to measure more than an upper bound on the strength of the bias. We allow the user to change these thresholds flexibly. Finally, if a database of known V genes for the species under consideration is provided, `grmlin` annotates the relationships of the alleles in the constructed germline database to already known V genes by aligning to all known V genes using `blastn v2.7.1` [1].

### Validation in human datasets

We apply our approach to bulk heavy chain repertoire sequencing of B cells from the peripheral blood of 11 pairs of twins (2 fraternal, and 9 identical), previously published in Ref. [7]. We reasoned that, since identical twins share their germline genotype, we should expect a high degree of concordance between their germline databases. Applied to this dataset, `grmlin` detects between 46 and 55 germline IGHV alleles in all individuals, with notable co-variation between pairs of twins (**Fig. 3a**). Though it does not rely on any database of known alleles, over 1000 of the genes called ‘germline’ by `grmlin` are exact matches to alleles found in the IMGT database of human IGHV alleles, 4% are found to deviate from a known allele by a single nucleotide, and a handful are found at further distances (**Fig. 3b**). Moreover, we detect a high degree of concordance between the germline databases of the twins (**Fig. 3c**): with the exception of one of the fraternal pairs (twins 1A/B), pairs of twins have remarkably consistent germline allele calls, which typically differ by no more than one detected allele. This should be contrasted with pairs of unrelated individuals in the same dataset, who share only about half of their alleles. Intriguingly, one of the fraternal pairs (twins 6A/B) exhibited a high degree of concordance consistent with the identical pairs. We emphasize that the concordance between pairs of alleles is not limited to alleles found in the IMGT database. In cases in which a novel allele is called as ‘germline’ in one of the individuals, the same allele is also independently called in their twin in 85% of the instances (**Fig. S1**).

**Figure 3.**
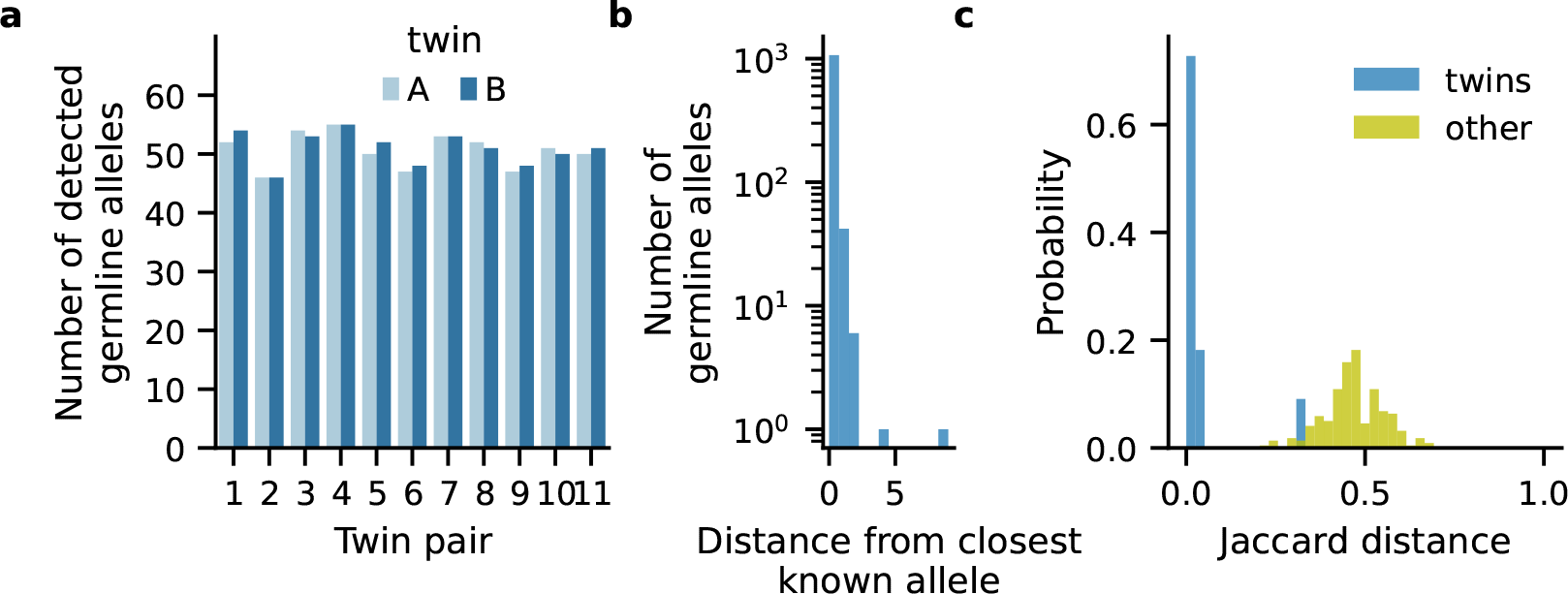
Properties of grmlin germline IGHV databases in 11 pairs of twins. (**a**) Number of detected germline IGHV alleles in each twin. (**b**) Hamming distances of all called germline IGHV alleles from closest gene in database. (**c**) Distribution of Jaccard distances between the germline IGHV databases of pairs of twins (blue) and pairs of unrelated individuals (light green).

We also used `partis`, `TIgGER` and `IgDiscover` to infer the germlines for the same individuals. Before providing the heavy chain transcript sequences to these tools, we used the same pre-processing procedure as for `grmlin`. We ran the all tools with default settings, except in the case with `IgDiscover`, where we lowered the `preprocessing_filter v_coverage` to 80, to generously accommodate for the V primer design of the twin dataset. Moreover, we used both the frequency-based and Bayesian implementations of `TIgGER`. Overall, we find that the frequency-based `TIgGER` and `partis` methods detect a comparable number of genes to `grmlin` but with poorer concordance between twins both in the full set of called IGHV germline genes (**Fig. 4**), and novel IGHV germline genes (**Fig. S1**). The Bayesian implementation of `TIgGER` detects far more alleles, but at a significant cost in similarity between pairs of twins (see **Fig. S1**). Finally, `IgDiscover` appears significantly less sensitive, detecting only between 8 and 13 germline V genes across all individuals.

**Figure 4.**
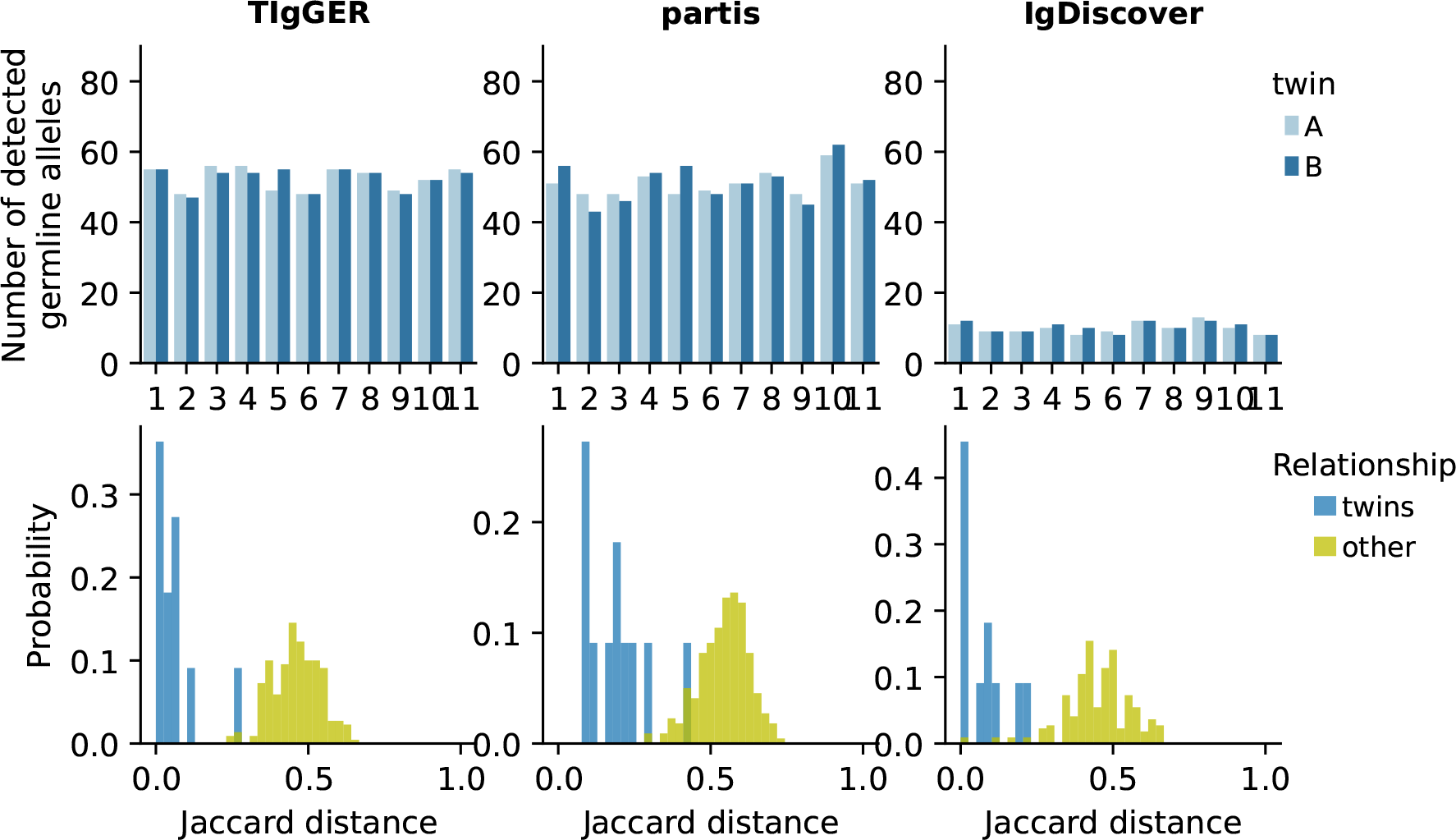
Comparison with germline IGHV databases constructed using IgDiscover and TIgGER. For each method, top panel shows the number of detected germline IGHV alleles in each twin, and the bottom panel shows the distribution of Jaccard distance between the germline IGHV databases of pairs of twins (blue) and pairs of unrelated individuals (light green).

## Discussion

Analysis of BCR repertoires often relies on the knowledge of the germline gene variants ancestral to the sampled BCR transcripts. This set of genes is highly personalized; even fraternal twins may share only about half of their germline IGHV genes. Thus, a variety of tools have been built to infer germline immunoglobulin genes from deep BCR sequencing data [2, 5, 12]. In this work, we presented a novel and orthogonal approach that achieves the same task without reference to a known reference database for a species. Our approach achieves this by making use of two empirical properties of B cell repertoires. First, naive B cells are highly abundant in B cell populations: thus the germline versions of Ig genes are present at high frequency in any dataset containing a high fraction of naive B cells. Second, hypermutated BCRs contain enormous sequence diversity, and convergent hypermutation of templated portions of V genes is rare. We built a simple software package that exploits these two properties, and we showed that it can achieve similar sensitivity and better specificity than previous methods, as measured by the concordance between pairs of twin germlines. We hope that our approach will be broadly useful to scientists studying both human BCR repertoires, as well of those of non-model organisms, as the empirical properties it exploits are likely to be universal.

Similarly to previous authors, we have implemented our approach with a primary focus on germline V genes, which harbor the most genetic diversity. We have found that our approach generally performs poorly in distinguishing germline variation from hypermutation in templated regions, and so it is unlikely to perform well on Ig D genes. A similar approach can in principle be implemented for Ig J genes. We have explored this, and found that it produces satisfactory results in templated regions, but often fails to distinguish human IGHJ4 and IGHJ5 genes, which differ primarily the presence of an additional codon on the 5’ of the IGHJ5 gene that is always a part of the CDR3 following VDJ recombination. We did not pursue a further implementation of personalized J gene databases in humans, especially because human IGHJ genes are generally not as diverse and appear well-annotated. In non-model organisms however, we anticipate that a similar approach may yield a good understanding of the templated portions of Ig J germline genes (i.e. outside the CDR3), with the important caveat that there may be additional sequence variation on the 5’ end of J genes.

A further limitation of our approach is that, because it relies on an abundance of naive B cells, it is fundamentally unsuited for use in datasets that consist of sorted mature B cell populations from which naive B cells have been excluded. Thus, we advise researchers to consider exploring alternative approaches if excluding naive B cells is critical to their experimental design, or including a bulk naive B cell library in their experimental design to facilitate simpler germline allele detection.

## Supplementary Information

**Figure S1.**
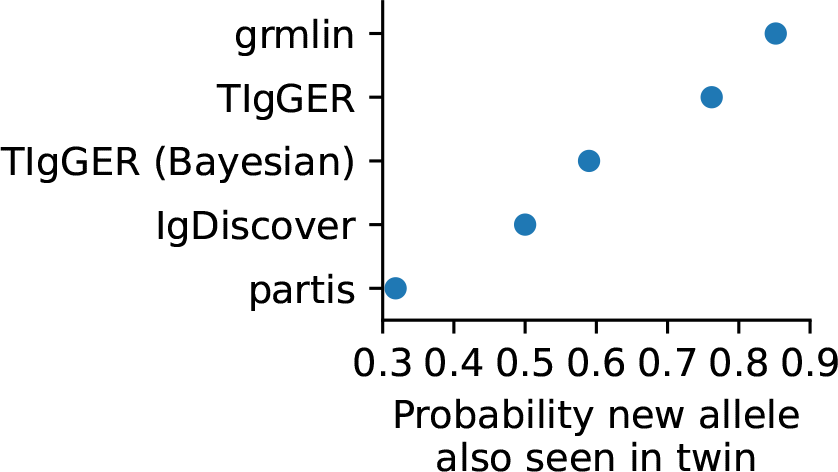
Probability that a novel allele is also detected in the twin, in each of the examined methods. We use ‘novel’ to refer to a germline allele without a perfect match in the IMGT database.

### S1 Probability of convergent hypermutation

In this section, we present a simply model of recurrent hypermutation, which we use to quantify the probability that two independently generated templated sequences that are an equal distance from their germline variant are identical, and show that the baseline rate at which these events arise is low.

**Figure S2.**
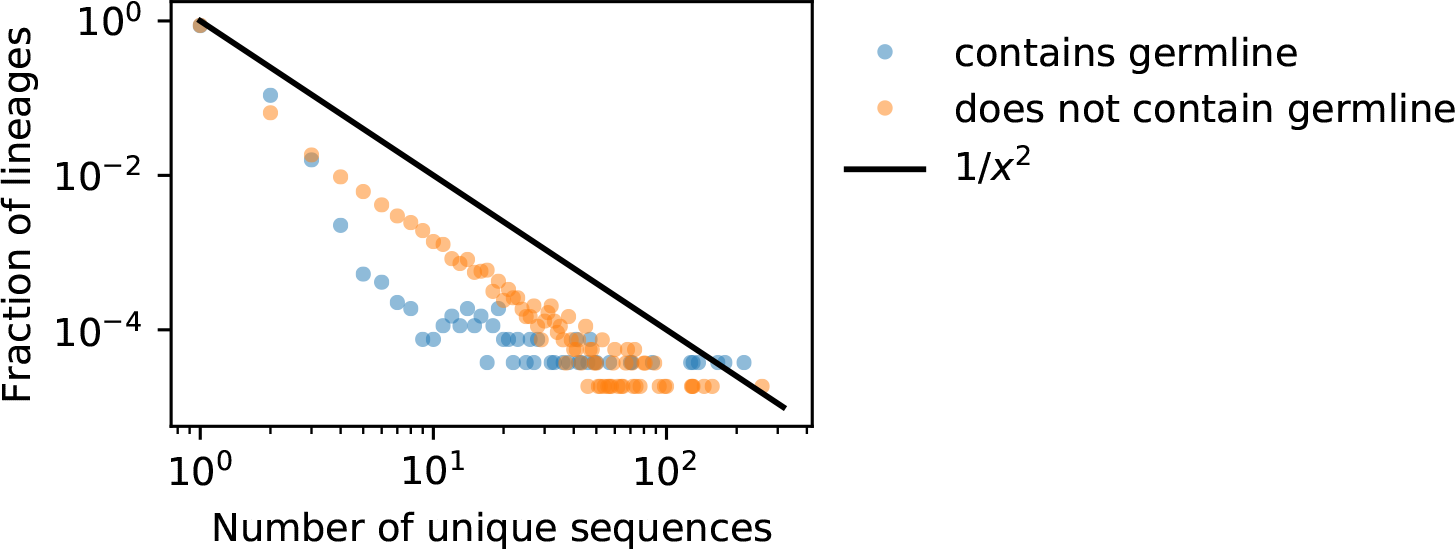
Distribution of the number of unique sequences in a lineage. Lineages were assigned on the basis of CDR3 Hamming distances, by single-linkage clustering of all sequences with CDR3s of equal length, the same IGHV family (as determined by IgBLAST) and 90% CDR3 sequence similarity. Blue points represent lineages that contain BCRs with IGHV sequences identical to those in the inferred germline database for this individual, and orange points represent lineages in which all BCRs contain hypermutations.

We will denote with *u*_*i,b*_ the rate at which a templated B cell sequence containing a particular SNV (*i, b*) is generated (specified by its position *i* and mutated base *b*),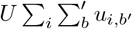 the sum of these rates, and with *μ*_*i*_ the fractional mutability (*μ*_*i*_ = *u*_*i*_*/U*). In independent lineages, the probabilities of convergent hypermutation are determined by these fractional mutabilities. The probability that that *n* independently generated V sequences that are a single mutation away from a germline segment share the same variant (*i, b*) is 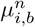 Similarly, the probability that *n* independently generated V sequences that are 2 mutations away from their ancestral germline segment all have the same SNVs (*i*_1_, *b*_1_) and (*i*_2_, *b*_2_) is equal to 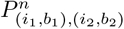 where

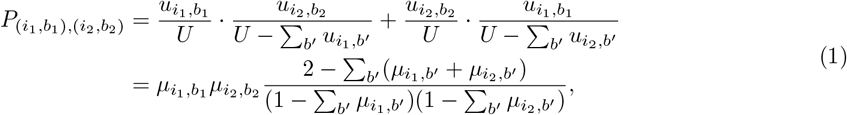

with analogous expressions for convergent hypermutation events a larger Hamming distance away from the germline.

When these fractional mutation rates are small compared to 1, these probabilities will be well approximated by

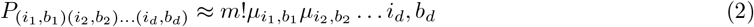

and the probability of hypermutating to any particular sequence a distance *d* apart from the germline will rapidly decline with *d*. In other words, convergently hypermutated sequences are expected to most readily arise in the vicinity of germline genes and their emergence is increasingly entropically suppressed as the distance from the germline increases.

## Acknowledgments

We thank Michael Swift for many helpful conversations while we were developing these ideas and for critical comments on the manuscript. IC acknowledges the support of the Lewis-Sigler Institute for Integrative Genomics and the Stanford Science Fellowship. ERJ acknowledges the Stanford University Chodorow Fellowship, and the Burroughs Wellcome Career Award at the Scientific Interface. Some of the computing for this project was performed on the Sherlock cluster. We would like to thank Stanford University and the Stanford Research Computing Center for providing computational resources and support that contributed to these research results.

---

**Figure S3.**
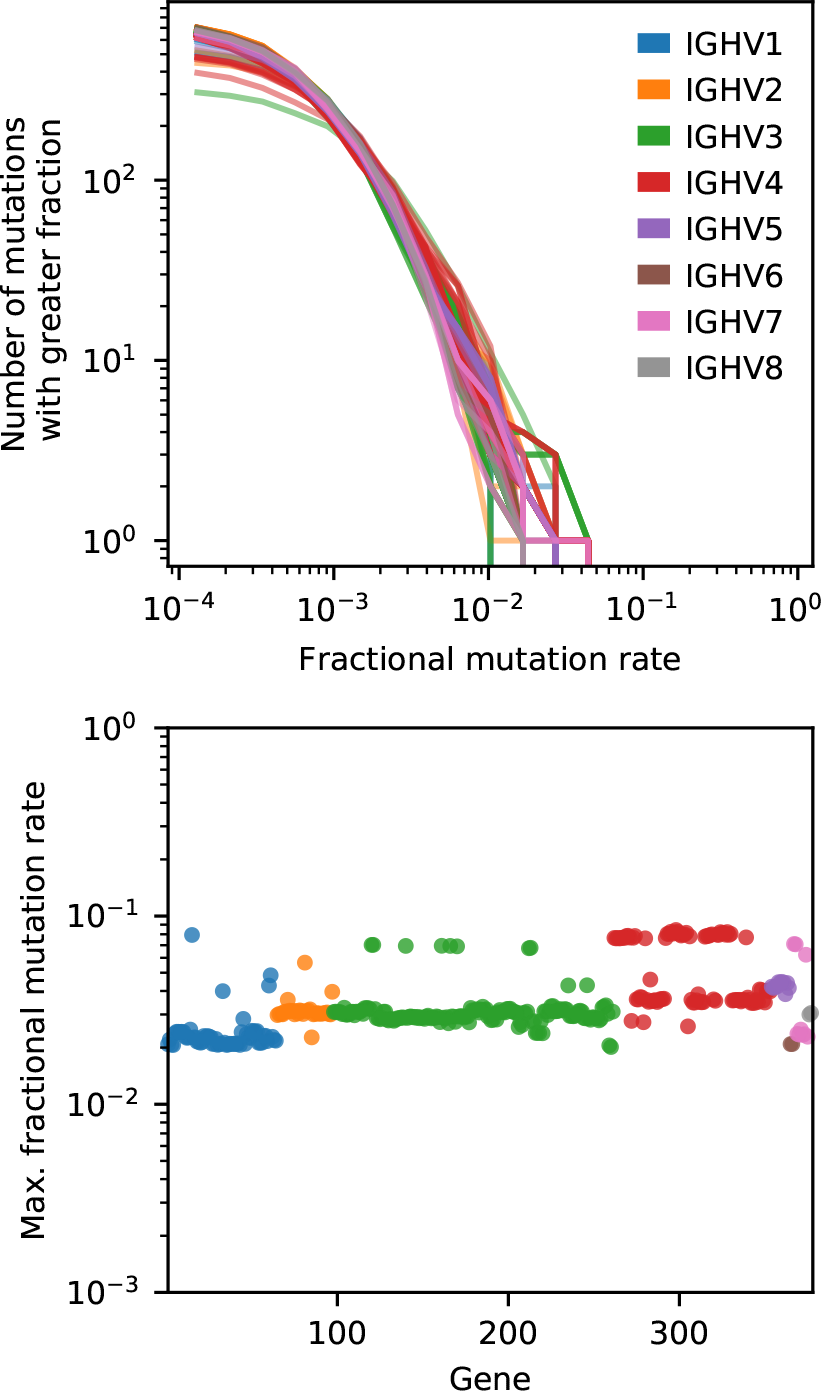
Fractional mutation rates at all positions for known IGHV genes. (Top panel) The distribution of fractional mutation rates of all possible SNVs for all IGHV genes in the IMGT database that do not contain an “N” in the reference allele, calculated using the motif and position-based ML phylogenetic model from unproductive sequences in Ref. [14]. (Bottom panel) The maximal fractional mutation rate for each gene.

**Figure S4.**
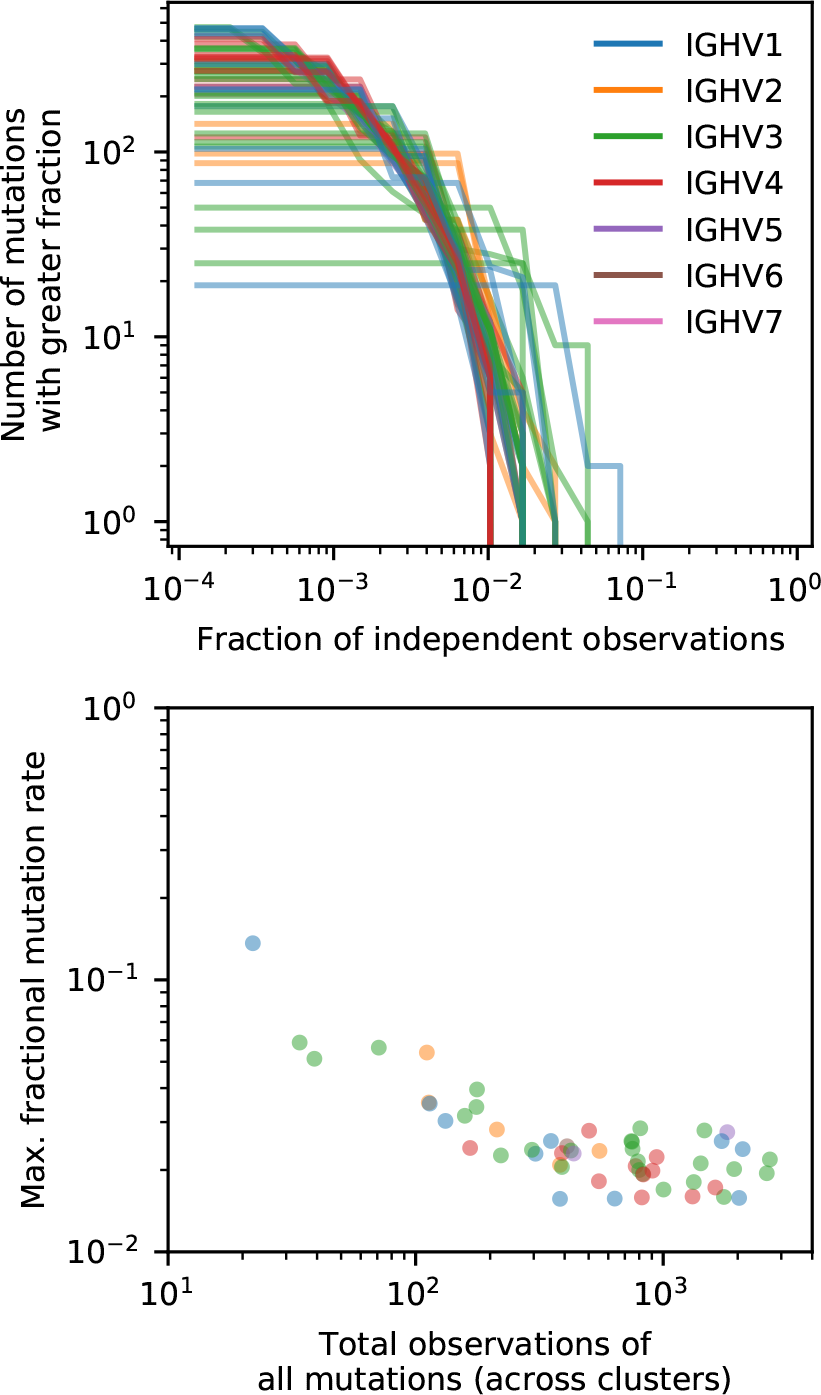
Estimates of relative mutation rates. Here, relative mutation rates are estimated by counting the number of multilineage CDR3 clusters (90% Hamming distance, single linkage) in which the mutation is observed. Each line (top panel) or dot (bottom) panel corresponds to one of the called germline genes in this sample (Twin 1B), which are colored by family. Mutations to different bases at the same site are counted separately.

## Notes

### Competing Interest Statement

The authors have declared no competing interest.

